# Inhibitory stabilized network behaviour in a balanced neural mass model of a cortical column

**DOI:** 10.1101/2022.12.09.519705

**Authors:** Parvin Zarei Eskikand, Artemio Soto-Breceda, Mark J. Cook, Anthony N. Burkitt, David B. Grayden

## Abstract

Strong inhibitory recurrent connections can reduce the tendency for a neural network to become unstable. This is known as inhibitory stabilization; networks that are unstable in the absence of strong inhibitory feedback because of their unstable excitatory recurrent connections are known as Inhibition Stabilized Networks (ISNs). One of the characteristics of ISNs is their “paradoxical response”, where perturbing the inhibitory neurons with additional excitatory input results in a decrease in their activity after a temporal delay instead of increasing their activity. Here, we develop a model of populations of neurons across different layers of cortex. Within each layer, there is one population of inhibitory neurons and one population of excitatory neurons. The connectivity weights across different populations in the model are derived from a synaptic physiology database provided by the Allen Institute. The model shows a gradient of excitation-inhibition balance across different layers in the cortex, where superficial layers are more inhibitory dominated compared to deeper layers. To investigate the presence of ISNs across different layers, we measured the membrane potentials of neural populations in the model after perturbing inhibitory populations. The results show that layer 2/3 in the model does not operate in the ISN regime but layers 4 and 5 do operate in the ISN regime. These results accord with neurophysiological findings that explored the presence of ISNs across different layers in the cortex. The results show that there may be a systematic macroscopic gradient of inhibitory stabilization across different layers in the cortex that depends on the level of excitation-inhibition balance, and that the strength of the paradoxical response increases as the model moves closer to bifurcation points.

**Author summary:** Strong feedback inhibition prevents neural networks from becoming unstable. Inhibition Stabilized Networks (ISNs) have strong inhibitory connections combined with high levels of unstable excitatory recurrent connections. In the absence of strong inhibitory feedback, ISNs become unstable. ISNs demonstrate a paradoxical effect: perturbing inhibitory neurons in an ISN by increasing their excitatory input results in a decrease in their activity after a temporal delay instead of increasing their activity. Here, we developed a neural mass model of a cortical column based on neurophysiological data. The model shows a gradual change in inhibitory stabilization across different layers in the cortex where layer 2/3 is less inhibitory stabilized and shows no paradoxical effect in contrast to layer 4 and layer 5, which operate in the ISN regime and show paradoxical responses to perturbation.

## Introduction

Even though cortical neurons receive a large number of feed-forward connections, mainly originating in the cortex, local recurrent connections heavily modulate their firing patterns [30, 31]. Neurophysiological data show that recurrent excitatory connections amplify feed-forward input signals, enabling pattern completion or other computations in the cortex, increasing the speed of network processing or increasing the capacity of memory storage [8, 12, 18, 19, 29]. Recurrent excitatory connections can lead to runaway excitation; however, strong feedback inhibition acts to prevent the network from becoming unstable [14, 27]. This is known as *inhibitory stabilization* and networks that combine high levels of excitatory recurrent connections with strong inhibitory feedback are known as Inhibition-Stabilized Networks (ISNs) [37].

Inhibitory stabilization plays an important role in maintaining balance in networks of cortical neurons. In the absence of stabilizing inhibitory feedback, the network may experience runaway activity resembling pathological states, such as epileptic seizures [20, 23]. However, impaired inhibitory stabilization does not necessarily result in catastrophic behavior such as seizures; in some cases, it may result in hyperactivity of the neurons or an increase in the cross-correlation between different types of neurons [36]. This highlights the importance of studying the detailed circuitry of balanced networks, the role of strong inhibitory connections in maintaining balance, and the effects of impaired inhibitory stabilization on the activity profiles of neurons.

One of the characteristics of ISNs is their *paradoxical response* where increasing the input to the inhibitory neural population results in a decrease of their activity instead of increasing their activity [32]. This is the signature of ISNs, and several experiments have been conducted to explore the possibility of the existence of ISNs in the cortex [1, 3, 16, 17, 33]. The presence of ISNs in the brain had been inferred through internal characteristics of the network of neurons, such as surround suppression or theta oscillations [28, 37]. Optogenetic stimulation experiments allow the possibility to investigate the direct effects of perturbing inhibitory neurons [6, 13, 40] and several experimental studies have shown evidence validating the presence of paradoxical effects in different areas of the cortex in awake mice [1, 16, 17, 33].

However, some experimental studies argue against the presence of ISNs in the cortex or show a mixture of results across different layers in the cortex [3, 21]. The study by Atallah et al.[3] showed no paradoxical effect in layer 2/3 of the primary visual cortex of anaesthetized mice and recent experimental work by Mahrach et al.[21] showed a paradoxical effect in layer 5 of the anterior lateral motor cortex (ALM) and barrel cortex but no paradoxical effect in layer 2/3 of the ALM.

In this study, a neural mass model (NMM) of a cortical column is developed based on parameters derived from recent anatomical and physiological studies by the Allen Institute [5, 34]. This data from the Synaptic Physiology database^1^ provides experimentally measured values of connectivity strengths across different populations of neurons in the mouse cortex. The connectivity weights of the NMM are determined based on this data and the resulting model is found to exhibit the differences in observations of the paradoxical effect across different layers in the cortex. The model shows that the presence of ISNs in the cortex is strongly tied to the level of excitation-inhibition balance. The model shows systematic variation of the connectivity strengths between populations that determine the gradient of excitatory and inhibitory balance across different layers in the cortex and, consequently, the degree of inhibitory stabilization. The difference in the level of excitatory and inhibitory balance across different layers in the cortex accords with neruophysiological data by Markram et al. [22] and Wang [39] who showed that superficial layers in the cortex have more inhibition-dominated activity compared to deeper layers, such as layer 5, that show more excitation-dominated connectivity.

## Methods

### The structure of the model

The model of the cortical column is constructed from three interconnected motifs, each of which is a model of a population of neurons (neural mass) in layer 2/3, 4 or 5, as illustrated in Fig. 1. Each motif contains a population model of excitatory neurons and a population model of inhibitory neurons. The population of excitatory neurons of each motif receives recurrent excitation from the excitatory population and inhibition from the motif’s inhibitory populations; the population of inhibitory neurons receives recurrent inhibition and excitation from the motif’s excitatory populations. The populations of neurons also receive excitatory and inhibitory input from other motifs in the network, as well as external excitatory inputs from outside of the column (thalamic and inter-cortical).

**Fig 1.**
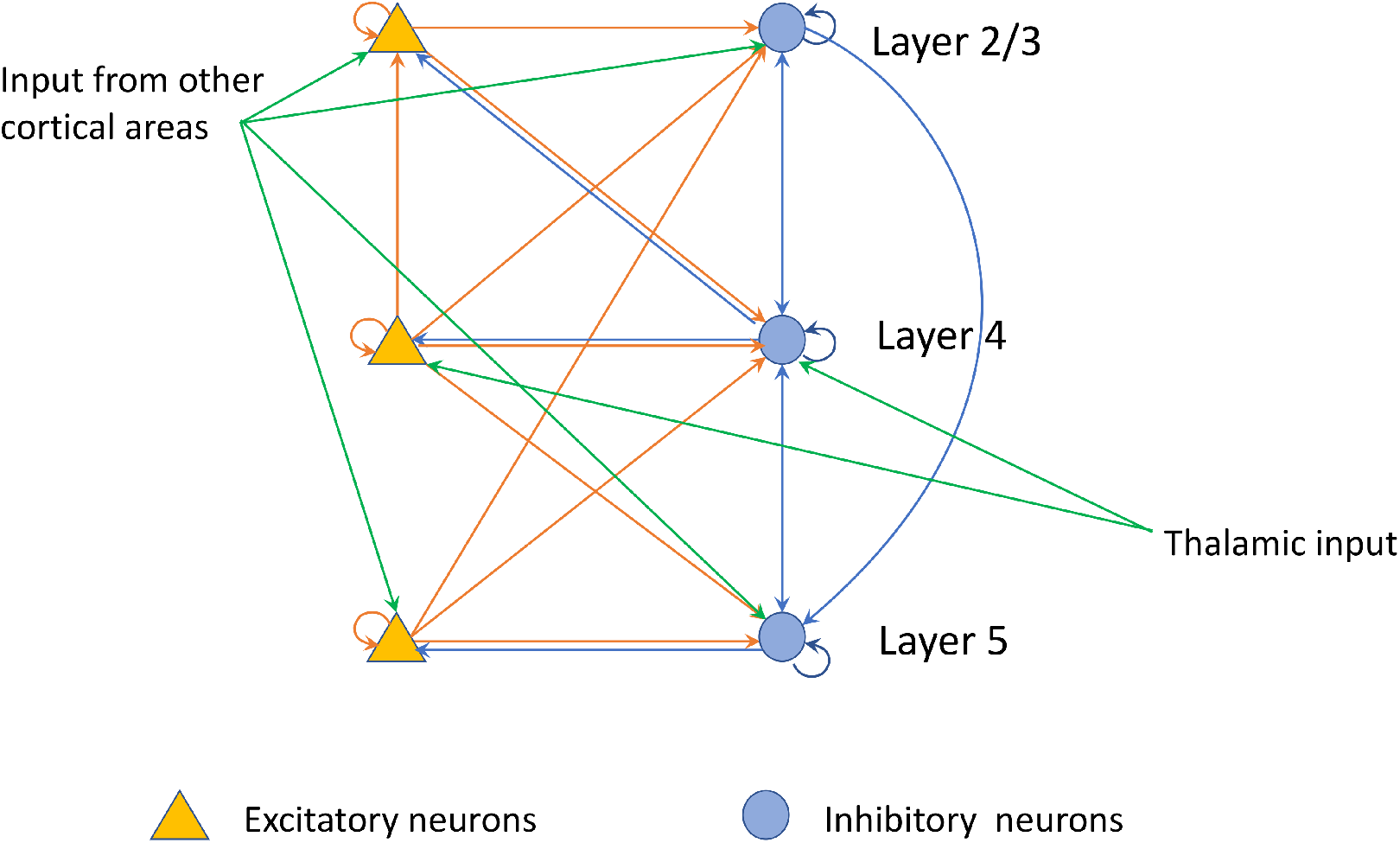
The structure of the neural mass model. The model contains populations of excitatory neurons and inhibitory neurons (motifs) in layers 2/3, 4 and 5 of the cortical column. The pattern and strengths of the connections between populations of neurons are set according to the values given in Tables 1 and 2. Yellow indicates excitatory populations and connections; blue indicates inhibitory populations of neurons and connections. Green arrows indicates the external excitatory input from thalamus and other cortical areas.

The pattern of connectivity between populations of neurons and the strengths of the excitatory and inhibitory connections are derived from the recently published Synaptic Physiology database (Allen Institute, USA) [5, 34]. This data provides estimates of the connection probabilities and synaptic strengths of the different classes of connections within a cortical column (see S1 Table and S2 Table, which are derived from Figure 4A and Figure 4B in [5]). The strengths and kinetics of the synapses in these experiments are measured by a double exponential fit to approximate the shape of post-synaptic potentials [34]. The results (reported in https://portal.brain-map.org/explore/models/mv1-all-layers) show that the rise-time constants vary between 0.6-1.9 ms across different types of inhibitory neurons and the decay time constants varies between 0.9-2.5 ms. The impact of the number of cells is taken into account under connection probability. Connectivity weights in our model are calculated by multiplying the connection probabilities *P_A,B_* and synaptic strengths *S_A,B_* for each pair of neural masses *A, B.* The database includes data from three types of inhibitory neurons: Htr3a, somatostatin (Sst) andparvalbumin (Pvalb); VIP neurons are considered to be a subclass of Htr3a in this database. The interconnections between different populations of the neurons in the model are based on the summation of the incoming synaptic currents. Therefore, for simplicity, we have considered all three types of neurons as one inhibitory population and added together their connectivity weights in the model. For example, the connectivity weights of inhibitory to excitatory connections in layer 2/3 is given by

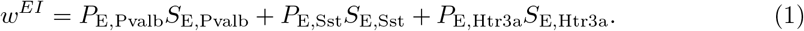

The connectivity weights for the intracolumnar connections are given in Table 1. All of the inter-layer connections for each motif are included in the model and for simplicity we have excluded weak intra-layer connections with connectivity weights below 0.1. The populations of neurons in layer 4 additionally receive thalamic input and the populations of neurons in layers 2/3 and 5 receive inputs representing other cortical areas, given in Table 2.

**Table 1.** Connectivity weights. The connectivity weights between populations of neurons in the model. E represents populations of excitatory neurons and I represents populations of inhibitory neurons in layers 2/3, 4 and 5. The highlighting indicates the strength of the connections, darker for the stronger connections.

|  |  |  | Postsynaptic |  |  |  |  |  |
| --- | --- | --- | --- | --- | --- | --- | --- | --- |
|  |  |  | Layer 2/3 |  | Layer 4 |  | Layer 5 |  |
|  |  |  | E2/3 | I2/3 | E4 | I4 | E5 | I5 |
| Presynaptic | Layer 2/3 | E2/3 | 0.06 | 0.88 | 0.01 | 0.25 | 0.06 | 0.16 |
|  |  | I2/3 | 0.35 | 1.06 | 0.06 | 0.18 | 0.02 | 0.06 |
|  | Layer 4 | E4 | 0.11 | 0.21 | 0.2 | 1.14 | 0.07 | 0.24 |
|  |  | I4 | 0.29 | 0.59 | 0.48 | 1.46 | 0.07 | 0.17 |
|  | Layer 5 | E5 | 0.01 | 0.13 | 0 | 0.13 | 0.09 | 0.27 |
|  |  | I5 | 0.04 | 0.06 | 0.02 | 0.11 | 0.49 | 1.51 |

**Table 2.** Parameters of the model.

| Parameter | Description | Value |
| --- | --- | --- |
| $\tau_m^E$ | Membrane time constant of the excitatory neurons | 0.02s [26] |
| $\tau_m^I$ | Membrane time constant of the inhibitory neurons | 0.04s [26] |
| $c$ | Membrane capacitance | 0.35nF [10] |
| $\tau_s^{EE}$ | Synaptic time constant for excitatory-to-excitatory synapses | 8.5 ms [2, 5] |
| $\tau_s^{IE}$ | Synaptic time constant for excitatory-to-inhibitory synapses | 4.3 ms [2, 5] |
| $\tau_s^{EI}$ | Synaptic time constant for inhibitory-to-excitatory synapses | 13 ms [2, 5] |
| $\tau_s^{II}$ | Synaptic time constant for inhibitory-to-inhibitory synapses | 9 ms [2, 5] |
| $r$ | Gain (steepness) of the sigmoid function | 0.3mV<sup>-1</sup> |
| $v_0$ | Postsynaptic potential for half of the maximum firing rate | 6 mV |
| $k$ | Connectivity constant | $5 \times 10^{-5}$ |
| $\phi_{\max}^E$ | maximum firing rate of excitatory neurons (where neurons saturate) | 21 Hz (inferred from [2]) |
| $\phi_{\max}^I$ | maximum firing rate of inhibitory neurons (where neurons saturate) | 30 Hz (inferred from [2]) |
| $w^E$ | Connectivity weight from other cortical areas/thalamus to excitatory populations | 0.1 (inferred from [25]) |
| $w^I$ | Connectivity weight from other cortical areas/thalamus to inhibitory populations | 0.2 (inferred from [25]) |

Each neural population is modeled as a rate-based unit with current-based synapses [7, 24]. The membrane potentials of the populations of neurons are given by the RC-circuit equation,

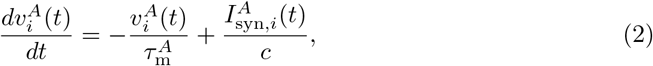

where 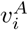 is the average membrane potential of a population of neurons, superscript A indicates either excitatory (*E*) or inhibitory populations (*I*), and the subscript *i* indicates the layer number (2/3, 4 or 5). The passive membrane time constants for each population *A*, 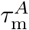, have different values for excitatory neurons and inhibitory neurons displayed in Table 2 [26]. The membrane capacitance is given by *c* = 0.35 nF. The differences in capacitance of different cell types result in different time constants and so are incorporated within the modelled time constants of the populations. 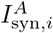 is the weighted summation of the incoming currents, 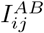, given by

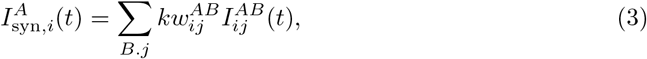

where 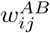 are the connectivity weights from population *B* in layer *j* to population A in layer *i,* displayed in Table 1, multiplied by a scaling connectivity constant, *k*. This scaling factor brings the values that are measured in single cell recordings to the correct operating regime for the neural mass models, while maintaining the relative strengths between them.

The dynamics of the post-synaptic currents are modelled using an exponentially decaying current [7, 9],

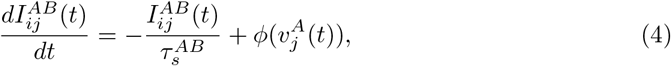

where 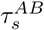 is the synaptic time constant associated with the particular synapse. For complete lists of equations of the model, see SI 1. The values of the synaptic time constants are derived from the recent computational study by Billeh et al. [5], who used alpha functions to describe synaptic currents based on electrophysiological studies by Arkhipov et al. [2]. Using MATLAB curve fitting tools (fit), we fit exponential functions to the alpha functions and calculated the resulting single-exponential synaptic time constants, which are given in Table 1 [2, 5].

The function *ϕ*() determines the output firing rate of a neural population by applying a sigmoid function to the membrane potential,

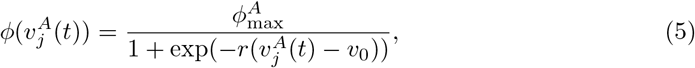

where 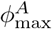 for A=E,I is the maximum firing rate of the neural population, *v*_0_ is the postsynaptic potential for which the firing rate reaches half of its maximum value and *r* determines the gain (steepness) of the sigmoidal firing rate function.We are making an approximation by choosing a universal output firing rate function for different populations.

Each population of neurons in the model receives input from other cortical areas or the thalamus. The dynamics of the synaptic equations for the input to excitatory and inhibitory populations, 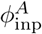 for A=E,I, follows the same patterns as other synaptic currents.

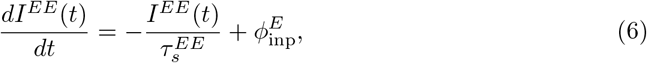

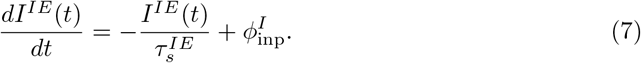

The default values for 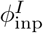 and 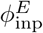 are 1 Hz which are chosen based on experimental data by Arkhipov et al. [2] showing the spontaneous firing rate of excitatory and inhibitory neurons are about 1 Hz.

### Excitation-inhibition balance

Excitation-inhibition balance is a fundamental characteristic of networks of neurons *in vivo* [15] and *in vitro* [35]. The excitation-inhibition balance in a layer of the model is evaluated with no driving external input and steady-state synaptic currents and membrane potentials. The balance value is calculated by subtracting the sum of the equilibrium synaptic inhibitory currents to the excitatory populations from the sum of the equilibrium excitatory currents to the excitatory populations [4, 38]. For example, the balance value, *B*, for layer 2/3 is defined as follows:

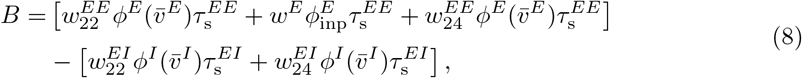

where 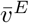 and 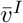 are the equilibrium membrane potentials of the excitatory and inhibitory populations, respectively.

### Fixed-point analysis

We evaluate the stability of the model by analysing the effects of small perturbations of the parameters. The first step is to find equilibria of the network by setting the left hand sides of the differential equations (i.e., all of the differential equations in the form of Eq. (2) and Eq. (4) equal to zero and solving the following equations:

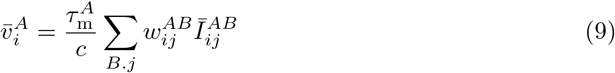

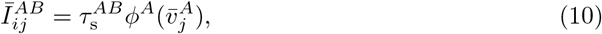

where 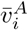 and 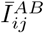 denote the equilibrium values of the variables. The complete detailed equations related to calculation of fixed points are in SI 2.

Bifurcation analyses are conducted using MatCont7p3 [11]^2^ to evaluate the behaviors of the network around the equilibria as the parameters of the model are changed. This is done for each layer (motif) separately and for the model as a whole.

### Paradoxical effect

To determine the existence of Inhibition-Stabilized Networks (ISNs) in each layer of the model, we examine the membrane potentials of the neural populations within each motif after perturbing its inhibitory population. The perturbation applied is a step increase in the firing rate of external excitatory input from 1 Hz to 4 Hz. We choose a substantial change in the firing rate of the perturbation to highlight the effect of perturbation on the membrane potential of the neural populations. We first analyze the motifs independently to remove effects of inter-layer interactions to look for ISN behavior. Then, we investigate the presence of ISNs in the whole cortical column by reintroducing excitatory and inhibitory interconnections across layers.

To measure the paradoxical effect, the membrane potential of the inhibitory populations is compared before and after applying the perturbation,

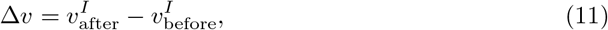

where 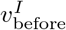 is the sustained (non-transient) membrane potential of the inhibitory population before perturbation and 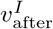 is the sustained membrane potential of inhibitory population after perturbation. A negative value of Δ*v* indicates a paradoxical response; i.e., the membrane potential reduces after the external input is increased. We evaluate changes in Δ*v* with variations in the parameters of the model.

## Results

### Excitation-inhibition balance

The balance in each motif between excitation and inhibition is evaluated separately for each layer of the model. Fig. 2 shows the balance values, B, with changing connectivity weights of inhibitory-to-inhibitory and inhibitory-to-excitatory connections between the neural populations within each motif. The default values for these connections are indicated by the stars. The other connectivity weights within each motif are set to the default values given in Table 1.

**Fig 2.**
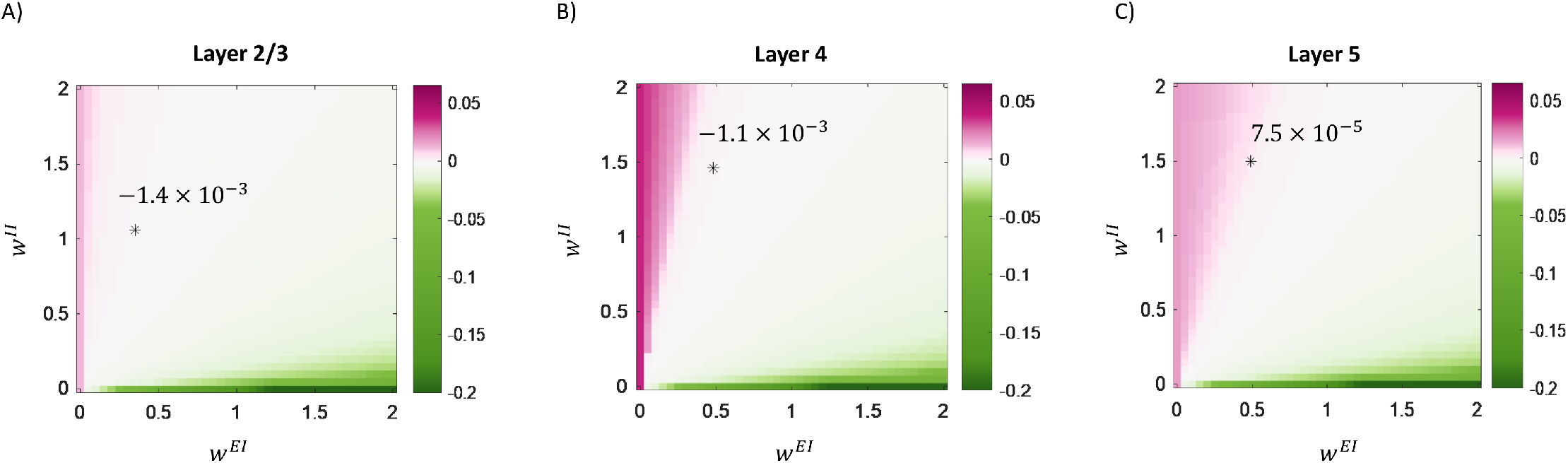
The balance level in each motif between excitatory and inhibitory populations of neurons. The colors indicate the balance levels in the individual motifs in the network for different values of inhibitory-to-excitatory and inhibitory-to-inhibitory connectivity weight between populations within (A) layer 2/3, (B) layer 4 and (C) layer 5. The x-axis shows the inhibitory-to-excitatory connectivity weight and the y-axis shows the inhibitory-to-inhibitory connectivity weight. The stars indicate the biological values of the two weights suggested by the Synaptic Physiology database. The balance value at the connectivity weights indicated by the star is displayed above the star.

The balance values at the starred points are very close to zero, showing near equal balance between excitation and inhibition in the motifs when the weights are set to the neurophysiologically derived values given in Table 2. The balance value in layer 5 is positive (7.5 × 10^-5^) compared to negative balance values of layer 2/3 (−1.4 × 10^-3^) and layer 4 (−1.1 × 10^-3^), showing that layer 5 is slightly more excitation dominated while the other layers are slightly more inhibition dominated. These results accord with the neurophysiological findings of [22], which showed that the connectivity in layer 5 of the somatosensory cortex of a juvenile rat is excitatory dominated compared to the superficial layers, such as layer 2/3.

### Fixed-point analysis of a single motif

We evaluate the stability of a motif when changing the connectivity weights. Fig. 3A shows the fixed-points analysis and changes in the firing rate of the excitatory neural population in a single motif as the weights are varied over the range [0, 2]. The colors indicate the firing rate of the neural population, which varied over the range [0, 21] for excitatory populations. Typically, the firing rates were close to zero (blue shades) or saturated (red shade).

**Fig 3.**
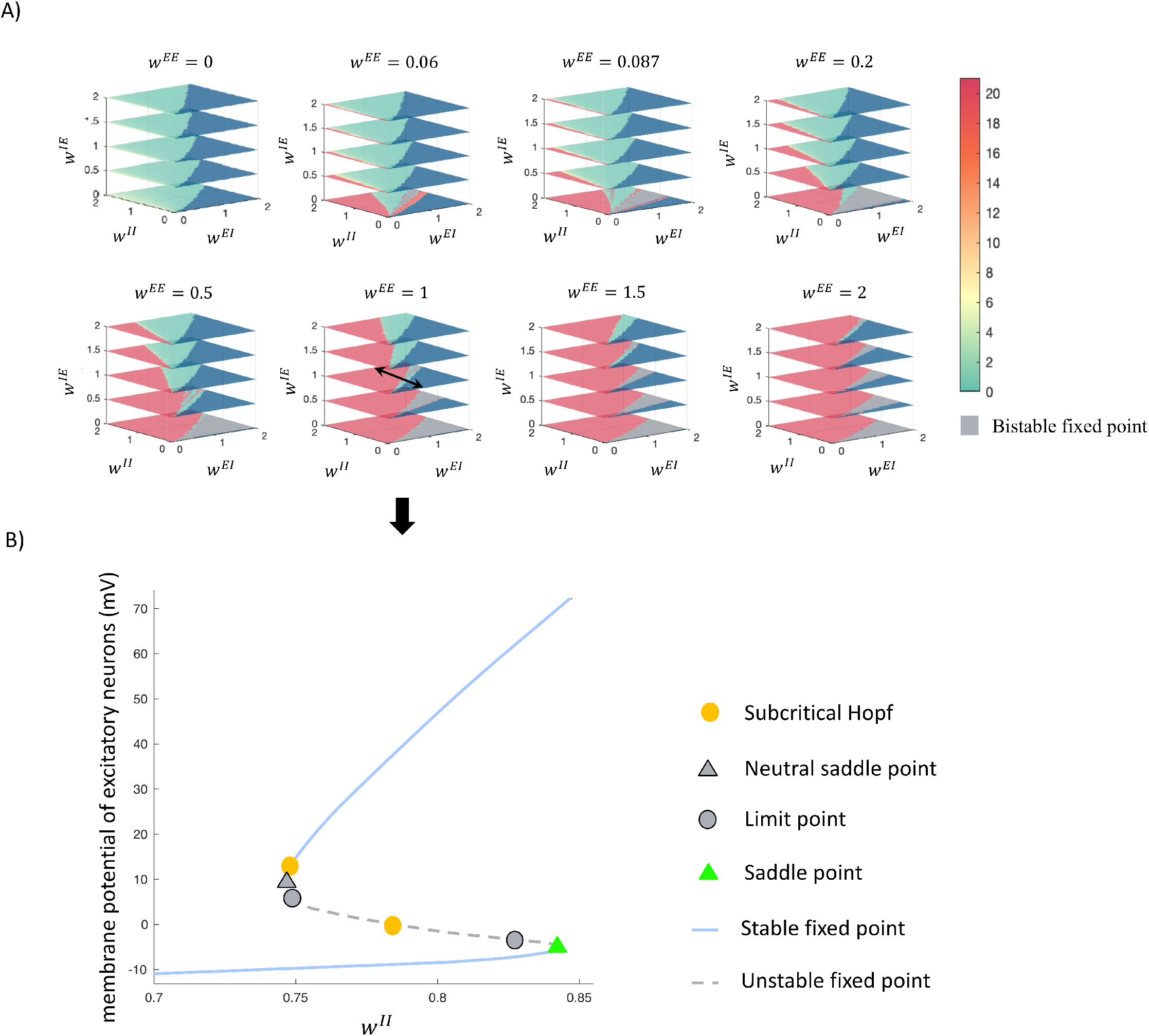
Fixed-point analysis of a single motif. A) Each block represents the firing rate of the excitatory neural population for different values of the network weights. The excitatory-to-excitatory connectivity weight increases from left to the right blocks as indicated above each plot. The axes shows different values of the connectivity weights for inhibitory-to-inhibitory, excitatory-to-inhibitory and inhibitory-to-excitatory connections. The colors indicate the firing rates, where the red color shows 21 Hz firing rate indicating the saturation at the maximum firing rate. Dark blue shows nearly zero firing rate. The gray color indicates the regions where the network has two stable fixed points (bistable). B) The detailed bifurcation analysis of the network where all of the connectivity weights have the value of 1 except the inhibitory-to-inhibitory connection (indicated by two-sided black arrow in A), which is the bifurcation parameter. The results show that the network has a saddle point (indicated by green triangle) where the stability of the fixed point changes. The network also has two subcritical Hopf bifurcations indicated by orange circles.

The grey regions represent values of the connectivity weights where the fixed points of the motif are in a bistable regime, where two stable states are separated by an unstable steady state (e.g., Fig. 3B). When the motif is in a bistable regime, a perturbation may cause the activity to move from one stable state to the other.

The detailed bifurcation analysis of the network indicates saddle-nodes or Hopf bifurcation at the border of the regions where the fixed points are bistable; i.e., indicated by a grey color in Fig. 3A. In these regions, a small perturbation generally causes the neurons to switch from low firing rate activity to high firing rates. Fig. 3B shows a detailed bifurcation analysis of one of the planes where the connectivity weights for all of the connections are equal to 1 except the inhibitory-to-inhibitory connectivity weights indicated by the two-sided black arrow in Fig. 3A. Changes in this parameter show two sub-critical Hopf bifurcation points (indicated by the orange circle). The analysis also shows a saddle point where the stability of the fixed point changes (indicated by the green triangle). In the regions where the fixed point is unstable, a small perturbation causes significant changes in the membrane potentials of the neurons from low activity to the saturation.

### Fixed-point analysis of the whole column

We also analyse the changes in the stability and firing rate of the neurons for different values of connectivity weights across the whole column model. Fig. 4 shows the results of this analysis when changing the connectivity weights of the inter-layer connections. All other connections have the default values indicated in Table 1.

**Fig 4.**
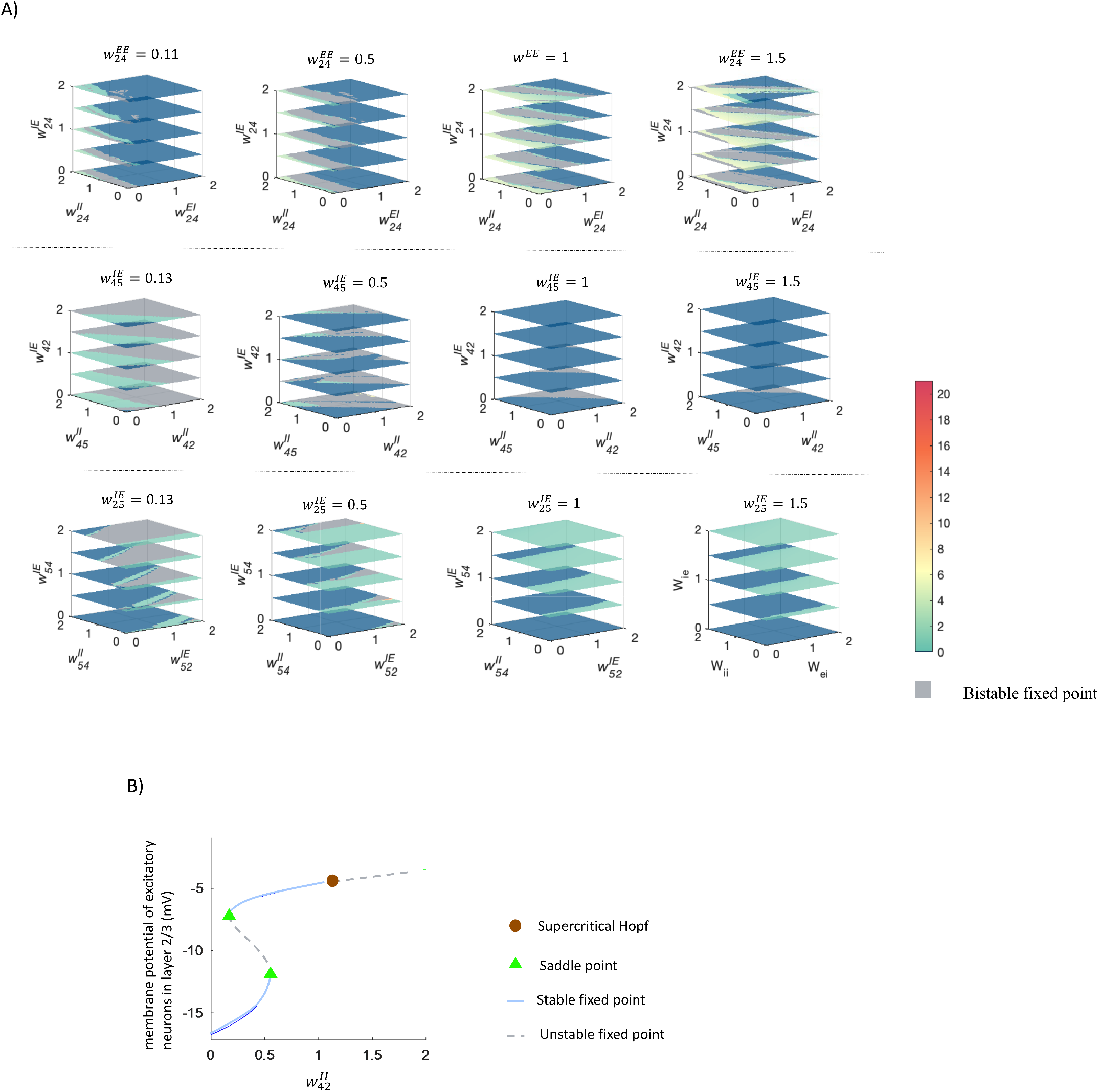
Fixed-point analysis of the whole column. A) Each block represents the firing rates of the excitatory neurons in layer 2/3 and the fixed-point analysis of the network for different values of connectivity weights. The axes and figure titles show different values of the connectivity weights of inter-layer connections. The colors indicate the firing rates of the excitatory neurons in layer 2/3, using the same scheme as Fig. 3. B) The detailed bifurcation analysis of the network where all of the connectivity weights have the default value set to the biological ranges in Synaptic Physiology database except the inhibitory-to-inhibitory connection from layer 2/3 to layer 4, which is the bifurcation parameter. The results shows that the network has two saddle points (indicated by green triangle) where the stability of the fixed point changes at this point. The network also has one supercritical Hopf bifurcations indicated by brown circle.

A detailed bifurcation analysis of the regions where the fixed-points are bistable, indicated by the gray color in Fig. 4A, reveals different behaviours of the network by changing the connectivity weights of inhibitory-to-inhibitory connections from layer 2/3 to layer 4 (second row in Fig. 4A). The bifurcation analysis indicates two saddle points and one super-critical Hopf bifurcation when changing the connectivity weights of inhibitory connections from layer 2/3 to layer 4 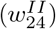 in a range of 0 to 2. As a result of this Hopf bifurcation, a stable limit cycle appears that generates an oscillations with a frequency of 12 Hz. The other regions with bistable fixed points in Fig. 4A (first and third rows) are the results of two saddle points, and varying the connectivity weights of these connections does not generate oscillatory behaviours.

### Layers 4 and 5 exhibit the paradoxical effect but layer 2/3 does not

We investigate the presence of Inhibitory-Stabilized Networks (ISNs) by looking for the paradoxical effect in each motif separately. Excitatory populations receive a default input of 1 Hz and inhibitory popoulations receive an input with a step increase perturbation at 1 second from 1 Hz to 4 Hz.

The layer 2/3 motif does not operate in the ISN regime: the membrane potential of inhibitory neural population increases to a higher steady-state value after the perturbation is applied. This is seen in Fig. 5A, which shows the membrane potentials of the inhibitory (blue) and excitatory (yellow) neural populations during the 2 s simulation period. After the input to the inhibitory population is increased, there is a transient period and then the membrane potentials settle on new values. The post-perturbation membrane potential of the inhibitory population is larger, showing the expected response when increasing its external stimulus.

**Fig 5.**
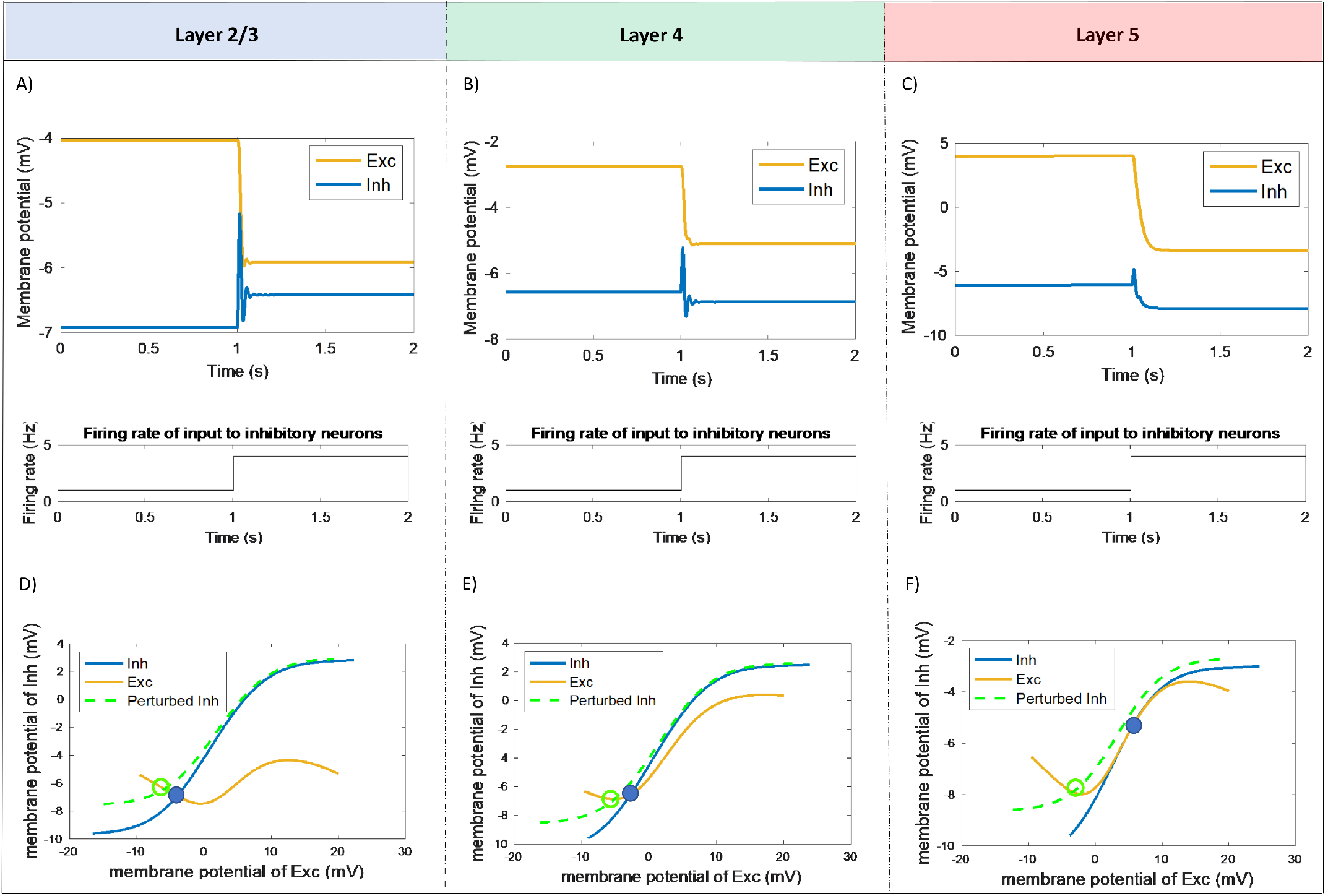
The paradoxical effect in individual motifs. A, B, C) Investigating the paradoxical effect across different isolated layers in the cortical column. The blue curve shows the membrane potential of inhibitory neurons and the orange curve shows the membrane potential of excitatory neurons. Layer 2/3 (A) does not show a paradoxical response. However, layer 4 (B) and layer 5 (C) show paradoxical responses to perturbation. D, E, F) The network dynamics and the location of the fixed point in layer 2/3 (D), layer 4 (E) and layer 5 (F). The x-axis is the membrane potential of excitatory neurons and y-axis is the membrane potential of inhibitory neurons. The blue curve shows the inhibitory nullcline and orange curve shows the excitatory nullcline. The inhibitory nullcline after perturbation is shown as the dashed green curve. The fixed point of the network before perturbation is shown by the blue filled circle and the fixed point after perturbation is shown by the green circle. The network shows the paradoxical effect when the fixed point of the network is located on the positive slope of the excitatory nullcline, which is the case for layer 4 (E) and layer 5 (F).

On the other hand, layers 4 and 5 operate in the ISN regime: the membrane potentials of the inhibitory neurons decrease after the perturbation to lower steady-state values (Figs. 5B and 5C).

Figs. 5D, E and F show the dynamics of each of the motifs. The derivatives of the membrane potentials of excitatory and inhibitory neural populations are set to zero to calculate the excitatory (yellow) and inhibitory (blue) nullclines. The point of intersection of the excitatory and inhibitory nullclines indicates the fixed point of the network. Where the fixed point of the network is located on the positive slope of the excitatory nullcline, we observe a paradoxical response after perturbation (the inhibitory nullcline shifts from the blue curve to the dashed green curve) as the location of the fixed point changes to a lower value of the membrane potential of inhibitory neurons as illustrated in Figs. 5E and F for layers 4 and 5, respectively. However, there is no paradoxical response when the fixed point of the network is located on the negative slope of the excitatory nullcline, as shown in Fig. 5D for layer 2/3.

Fig. 6 shows the effects of changes in the connectivity weights of excitatory-to-excitatory connections on the behaviours of the motifs. The first row (Figs. 6A, B, C) shows how Δ*v* (purple) and the firing rates (yellow for excitatory neurons, blue for inhibitory neurons) vary with excitatory-to-excitatory (EE) connectivity weights for each motif. The stars indicate the values of Δ*v* for the default weights. The paradoxical effect is present when Δ*v* is below zero (dashed green line). As the EE weight increases, the paradoxical effect becomes more pronounced until a critical point at which the firing rates of the excitatory neurons saturate, a point of failure of inhibitory stabilization.

**Fig 6.**
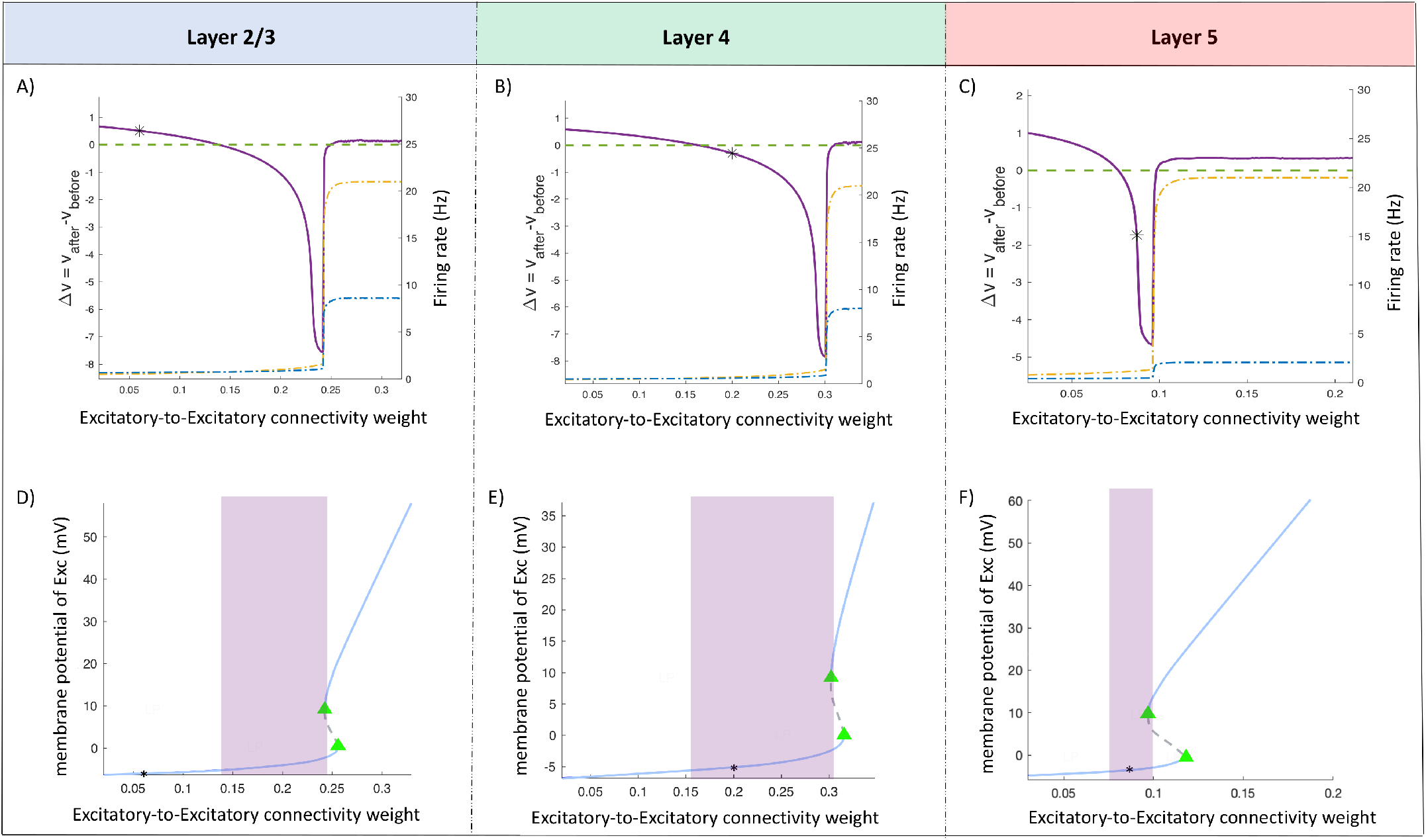
The effect of variation in the excitatory-to-excitatory connectivity weight on the paradoxical effect in single motifs. A, B, C) The purple curve indicates changes in the membrane potential (Δ*v*) of inhibitory neurons after perturbation for different excitatory-to-excitatory connectivity weights. The green dashed line indicates where Δ*v* is zero. Negative values of Δ*v* indicates paradoxical behavior. The black star indicates the default value of the excitatory-to-excitatory connectivity weight. The orange dashed line shows the firing rate (right axis) of excitatory population and the blue dashed line the firing rate of inhibitory population. Increasing the excitatory-to-excitatory weight from the default value results in the paradoxical effect in layer 2/3 (A) and increases the strength of the paradoxical response for all motifs. Excessive increase in the value of excitatory-to-excitatory connectivity weight results in saturation of the excitatory neurons. D, E, F) Bifurcation analysis of the motifs in layer 2/3 (D), layer 4 (E) and layer 5 (F). The y-axis is the membrane potential of the excitatory population and the x-axis is the bifurcation parameter, which is the excitatory-to-excitatory connectivity weight. The blue line indicates stable fixed points and the dashed gray line indicates unstable fixed points. The green triangles indicate saddle node bifurcations. The black star shows the default value of the excitatory-to-excitatory connectivity weight. The purple rectangle indicates the region where the motif operates in the ISN regime and shows a paradoxical response, which is obtained from the figures in the first row.

Figs. 6D, E, F show corresponding bifurcation analyses of the motifs with regard to the ISN regions (shown as purple shaded regions). Increasing EE weights results in gradually increasing membrane potential of the excitatory neural population until a saddle node bifurcation occurs (when *w^EE^* = 0.242 for layer 2/3 in Fig. 6D, *w^EE^* = 0.302 for layer 4 in Fig. 6E and *w^EE^* = 0.097 for layer 5 in Fig. 6F). Prior to this weight value in each case, the system reaches a point where there are multiple fixed points, and input perturbation causes the population to switch to the high membrane potential and subsequently high firing rate state. S1 Figure shows the variations in the paradoxical response and the location of the fixed point in the network by changing the excitatory-to-excitatory connectivity weight in layer 2/3.

Fig. 7A, B and C shows how Δ*v* (purple) and the firing rates (yellow for excitatory neurons, blue for inhibitory neurons) vary with excitatory-to-inhibitory (IE) connectivity weight for each motif. As the IE weight decreases, the paradoxical effect becomes more pronounced until a critical point at which the firing rate of the excitatory population saturates. The second row in Fig. 7 shows the bifurcation analyses when changing excitatory-to-inhibitory (IE) connectivity weights. Decreasing EI increases the strength of the paradoxical response to perturbation (negative magnitude of Δ*v*) (see Fig. 7A, B, C) until a saddle node bifurcation occurs similar to the behaviors seen in Fig. 6.

**Fig 7.**
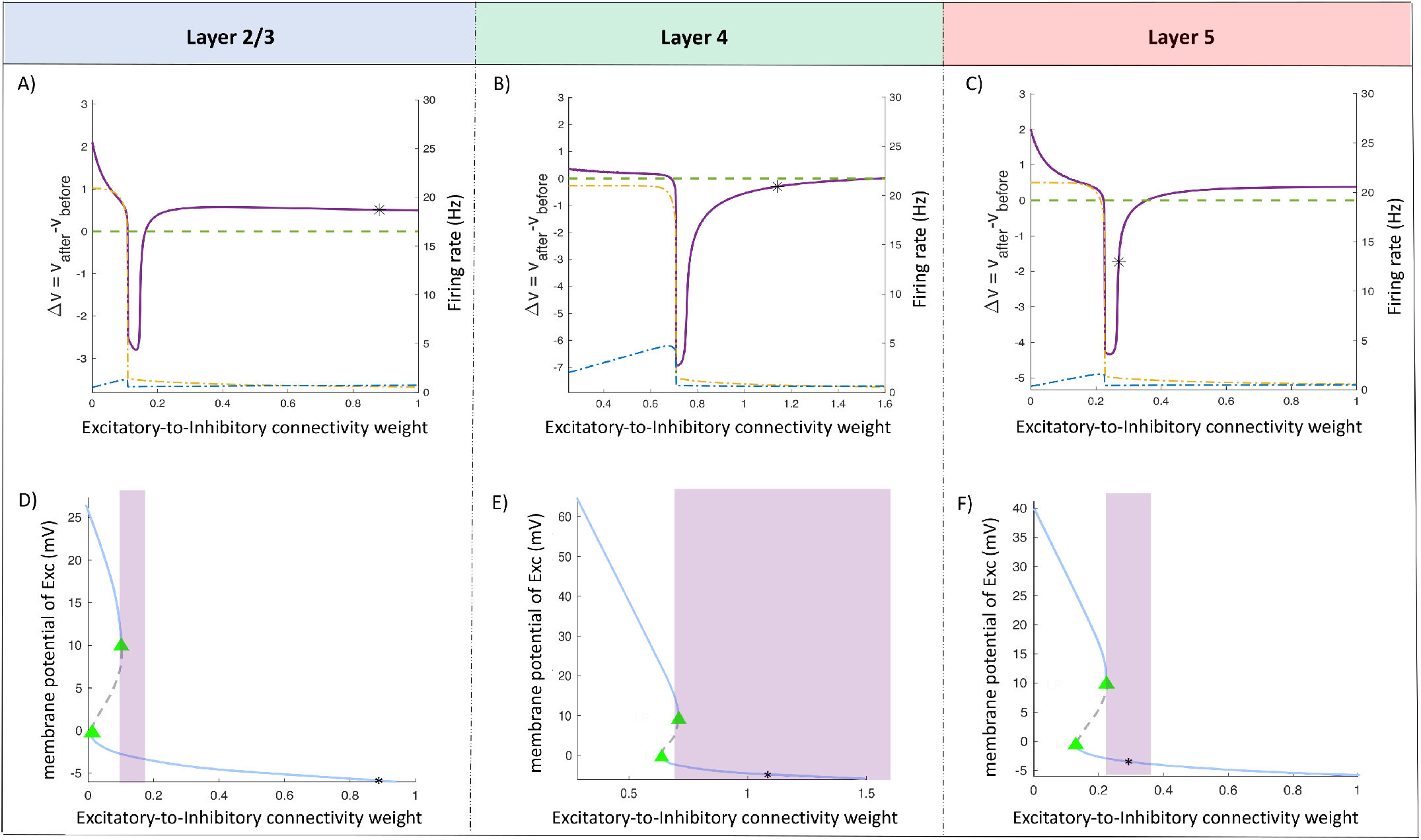
The effect of variation in the excitatory-to-inhibitory connectivity weight on the paradoxical effect in single motifs. A, B, C) The purple curve indicates changes in the membrane potential (Δ*v*) of inhibitory neurons after perturbation for different excitatory-to-inhibitory connectivity weights. The green dashed line indicates where Δ*v* is zero. Negative values of Δ*v* indicates paradoxical behaviour. The black star indicates the default value of the excitatory-to-inhibitory connectivity weight. The orange dashed line shows the firing rate (right axis) of excitatory population and the blue dashed line the firing rate of inhibitory populations. Decreasing the excitatory-to-inhibitory weight from the default value results in the paradoxical effect in layer 2/3 (A) and increases the strength of the paradoxical response in all motifs until a bifurcation occurs and the firing rate of excitatory neurons switches to 21 Hz. Excessive decrease in the value of excitatory-to-inhibitory connectivity weight results in saturation of the excitatory neurons. D, E, F) Bifurcation analysis of the motifs in layer 2/3 (D), layer 4 (E) and layer 5 (F) by variation in the excitatory-to-inhibitory connectivity weight. The y-axis is the membrane potential of the excitatory populations and x-axis is the bifurcation parameter, which is the excitatory-to-inhibitory connectivity weight. The blue line indicates stable fixed points and the dashed gray line indicates unstable fixed point. The green triangles indicate saddle node bifurcations. The black star shows the default value of the excitatory-to-inhibitory connectivity weight. The purple rectangle indicates the region where the motif operates in the ISN regime and shows a paradoxical response.

Fig. 8 shows the ranges of values for the inhibitory-to-excitatory connections where the motifs operate in the ISN regime. Decreasing inhibitory-to-excitatory connectivity weights leads to operation in the ISN regime with further decrease resulting in saturation of the excitatory populations, with corresponding bifurcation analyses similar to those in Fig. 7.

**Fig 8.**
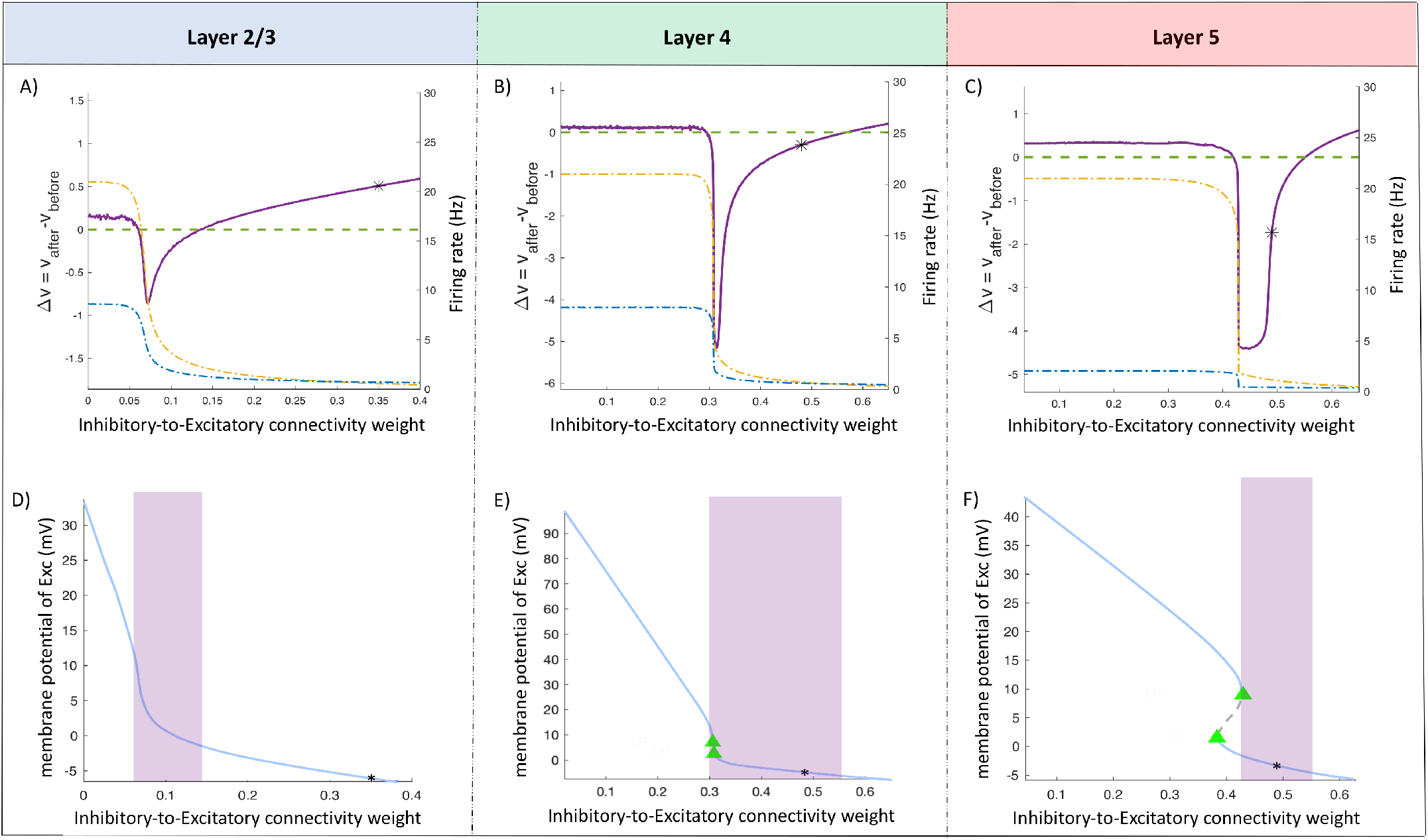
The effect of variation in the inhibitory-to-excitatory connectivity weight on the paradoxical effect in single motifs. A, B, C) The purple curve indicates changes in the membrane potential (Δ*v*) of inhibitory neurons after perturbation for different inhibitory-to-excitatory connectivity weight. The green dashed line indicates where Δ*v* is zero. Negative values of Δ*v* indicates paradoxical behaviour. The black star indicates the default value of the inhibitory-to-excitatory connectivity weight. The orange dashed line shows the firing rate (right axis) of excitatory population and the blue dashed line the firing rate of inhibitory population. Decreasing the inhibitory-to-excitatory from the default value results in the paradoxical effect in layer 2/3 (A) and increases the strength of the paradoxical response for all motifs. Excessive decrease in the value of inhibitory-to-excitatory connectivity weight results in saturation of the excitatory neurons. D, E, F) Bifurcation analysis of the motifs in layer 2/3 (D), layer 4 (E) and layer 5 (F). The y-axis is the membrane potential of the excitatory population and the x-axis is the bifurcation parameter, which is the inhibitory-to-excitatory connectivity weight. The blue line indicates stable fixed point and the dashed gray line indicates unstable fixed points. The green triangles indicate saddle node bifurcations. The black star shows the default value of the inhibitory-to-excitatory connectivity weight. The purple rectangle indicates the region where the motif operates in the ISN regime and shows a paradoxical effect.

Fig. 9 shows the ranges of values for the inhibitory-to-inhibitory connections where the motifs operate in the ISN regime. Increasing inhibitory-to-inhibitory connectivity weights leads to operation in the ISN regime. Further increase in this connectivity weight results in saturation of excitatory populations. In the absence of inhibitory-to-inhibitory connection (i.e., when the inhibitory-to-inhibitory connectivity weights equal zero), the excitatory populations stop firing and inhibitory populations saturate. The second row in Fig. 9 shows the bifurcation analyses when changing the inhibitory-to-inhibitory (II) connectivity weights in isolated motifs.

**Fig 9.**
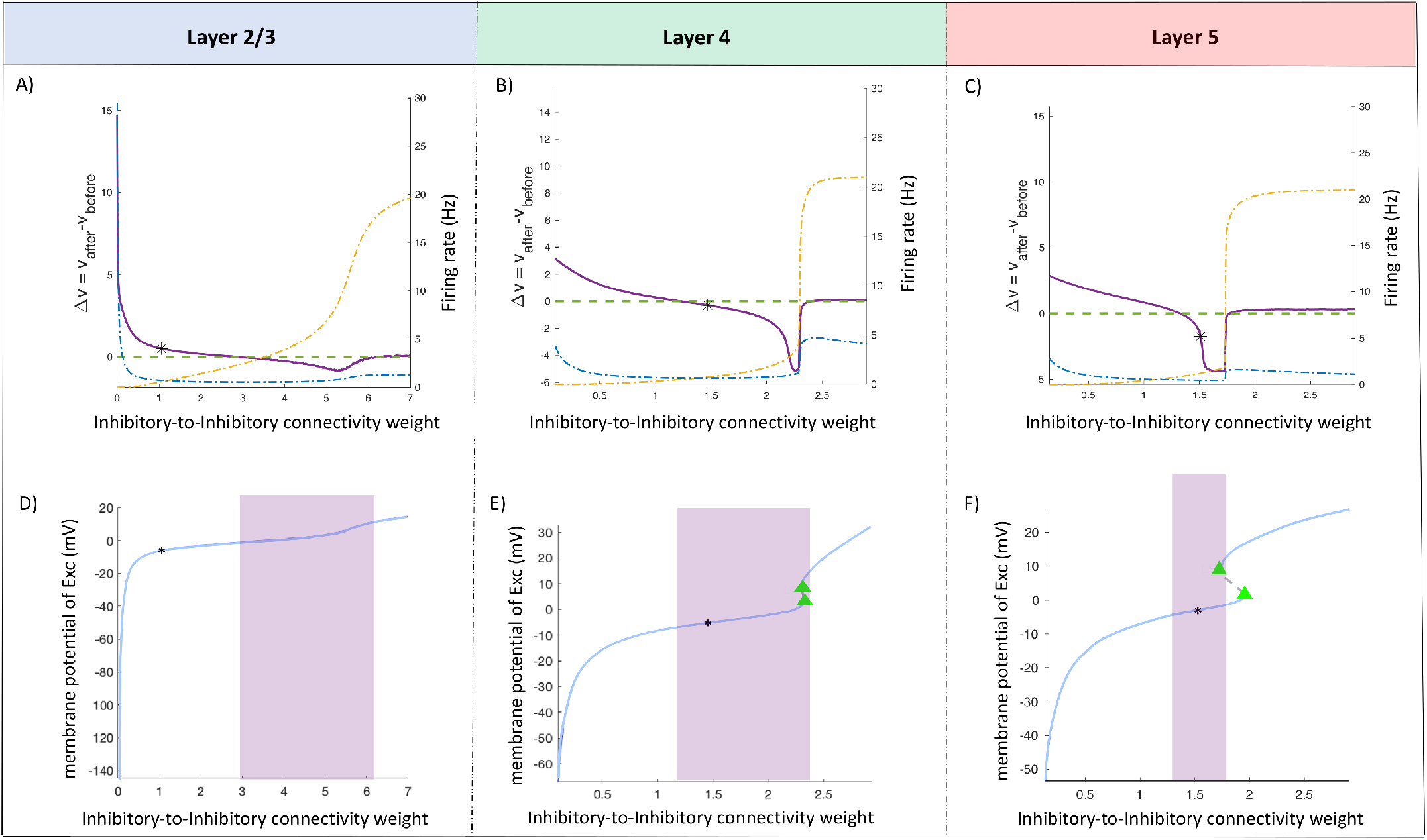
The effect of variation in the inhibitory-to-inhibitory connectivity weight on paradoxical effect in single motifs. A, B, C) The purple curve indicates changes in the membrane potential (Δ*v*) of inhibitory neurons after perturbation for different inhibitory-to-inhibitory connectivity weight. The green dashed line indicates where Δ*v* is zero. Negative values of Δ*v* indicates paradoxical behaviour. The black star indicates the default value of the inhibitory-to-inhibitory connectivity weight. The orange dashed line shows the firing rate (right axis) of excitatory populations and the blue dashed line the firing rate of inhibitory populations. Increasing the inhibitory-to-inhibitory connectivity weight from the default value results in the paradoxical effect in layer 2/3 (A) and increase the strength of the paradoxical response for all motifs. Excessive increase in the value of inhibitory-to-inhibitory results in saturation of the excitatory neurons. D, E, F) Bifurcation analysis of the single motif in layer 2/3 (D), layer 4 (E) and layer 5 (F). The y-axis is the membrane potential of the excitatory populations and the x-axis is the bifurcation parameter, which is the inhibitory-to-inhibitory connectivity weight. The blue line indicates stable fixed points and the dashed gray line indicates unstable fixed point. The green triangles indicate saddle node bifurcations. The black star shows the default value of the inhibitory-to-inhibitory connectivity weight. The purple rectangle indicates the region where the motif operates in the ISN regime and shows a paradoxical effect.

Results for each layer of the model when simulating the whole cortical column are shown in Fig. 10. Results were obtained for each layer by perturbing the populations of inhibitory neurons and recording from excitatory and inhibitory neurons of different layers. With the default parameter values, the column operates in the ISN regime in layer 4 (Fig. 10B, Δ*v* = −0.02) and layer 5 (Fig. 10C, Δ*v* = −0.9), but not layer 2/3 (Fig. 10A, Δ*v* = 0.09). These results show that the operation of the networks in layer 5 and layer 4 operate in ISN regime does not depend on the inhibitory connections from other layers. This is also the case for the network in layer 2/3. We observe paradoxical effect in layer 5 and layer 4 but not in layer 2/3 regardless of inhibitory connections across different layers in the cortical column.

**Fig 10.**
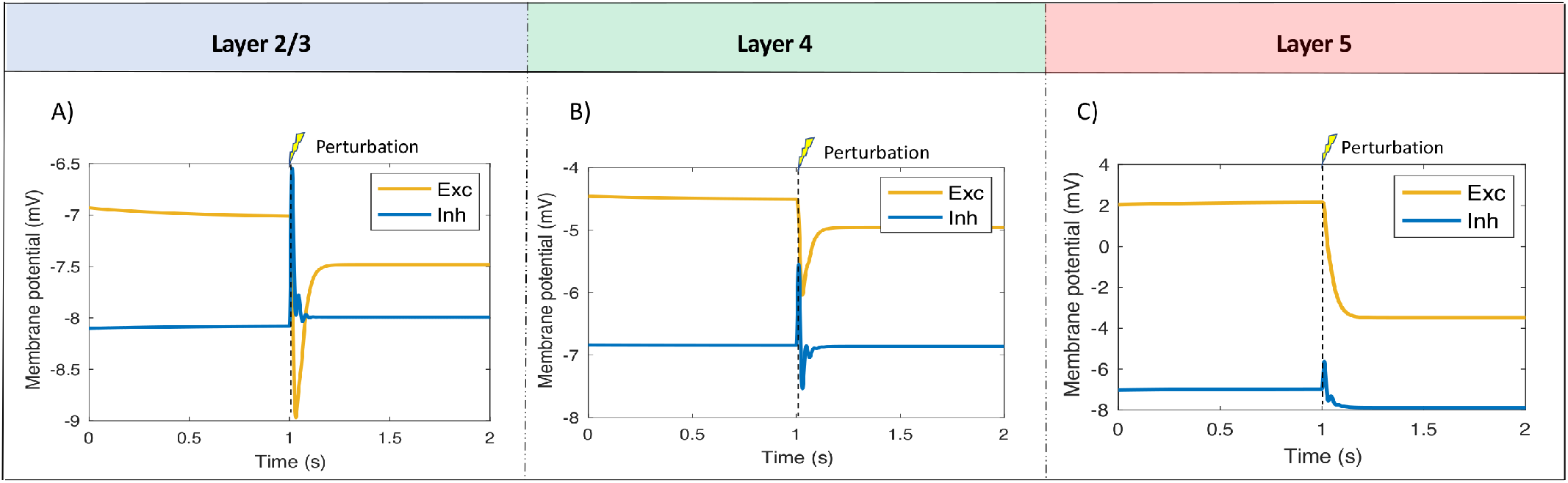
The paradoxical effect in the model of the whole column. The responses of populations in different layers of the cortical column. The membrane potential of excitatory populations are shown by the orange curves and the membrane potential of inhibitory populations are shown by the blue curves. Layer 2/3 (A) does not show the paradoxical effect. Layer 4 (B) and layer 5 (C) show paradoxical responses to perturbation.

### Evaluation of robustness of the model

The effects of varying the fixed parameters of the model were investigated to examine the robustness of the model to these parameters. These included the connectivity constant, *k,* and the saturation firing rates of the neural populations, 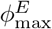 and 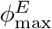 (see Table 2).

Fig. 11 shows the effects of varying the connectivity constant, *k*, on the paradoxical effect in single motifs when the connectivity weights are set to their default values. Negative values of Δv, shown by the purple curves, indicate values of *k* the paradoxical effect is present. The black stars indicate the default value of the connectivity constant used in the model. The results show that layer 2/3 does not show the paradoxical effect over a wide range of the connectivity constant (Fig. 11A). Layers 4 and 5 show increasing degrees of the paradoxical effect with increasing *k*, with smaller values resulting in non-paradoxical behaviour. Layer 5 shows the greatest sensitivity to *k,* with a large increase resulting in saturation of the excitatory neurons (orange dashed line in Fig. 11C).

**Fig 11.**
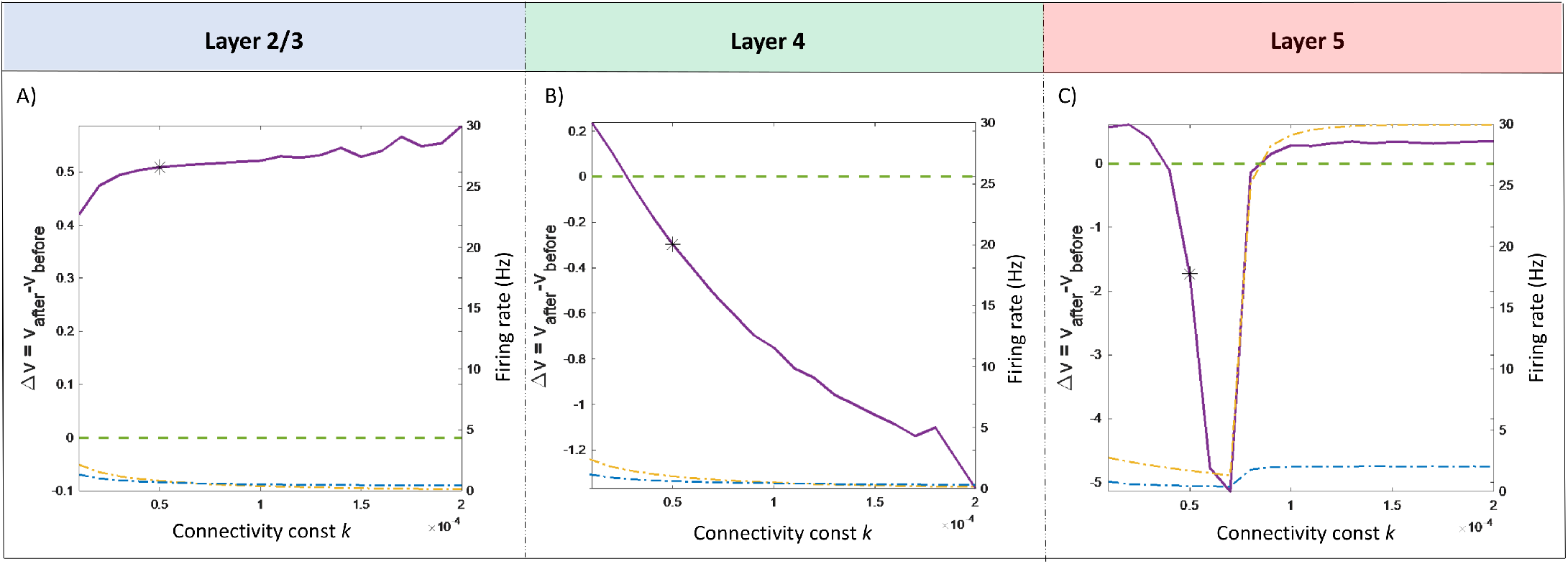
The effect of variation of the connectivity constant on the paradoxical effect in single motifs. A, B, C) The purple curve indicates changes in the membrane potential (Δ*v*) of inhibitory neurons with perturbation for different values of the connectivity constant. The green dashed line indicates where Δ*v* is zero. Negative values of Δ*v* indicate paradoxical behaviour. The black star indicates the default value of the connectivity constant. The orange dashed line shows the firing rate (right axis) of the excitatory population and the blue dashed line the firing rate of the inhibitory population.

Figs. 12A, B, C show the effects on the paradoxical effect in single motifs of varying the level of external input using the default sigmoid function. Layer 2/3 does not show the paradoxical effect for a wide range of the input levels except when the input is above 9 Hz. Layers 4 shows the paradoxical effect regardless of the input level. Layer 5 operates in the ISN regime over a narrow range of input values, with higher inputs leading to saturation and lower inputs reducing or eliminating the paradoxical effect. The default value of the external input was chosen as it generates firing rates close to 1 Hz, the observed spontaneous firing rates of neurons in cortex.

**Fig 12.**
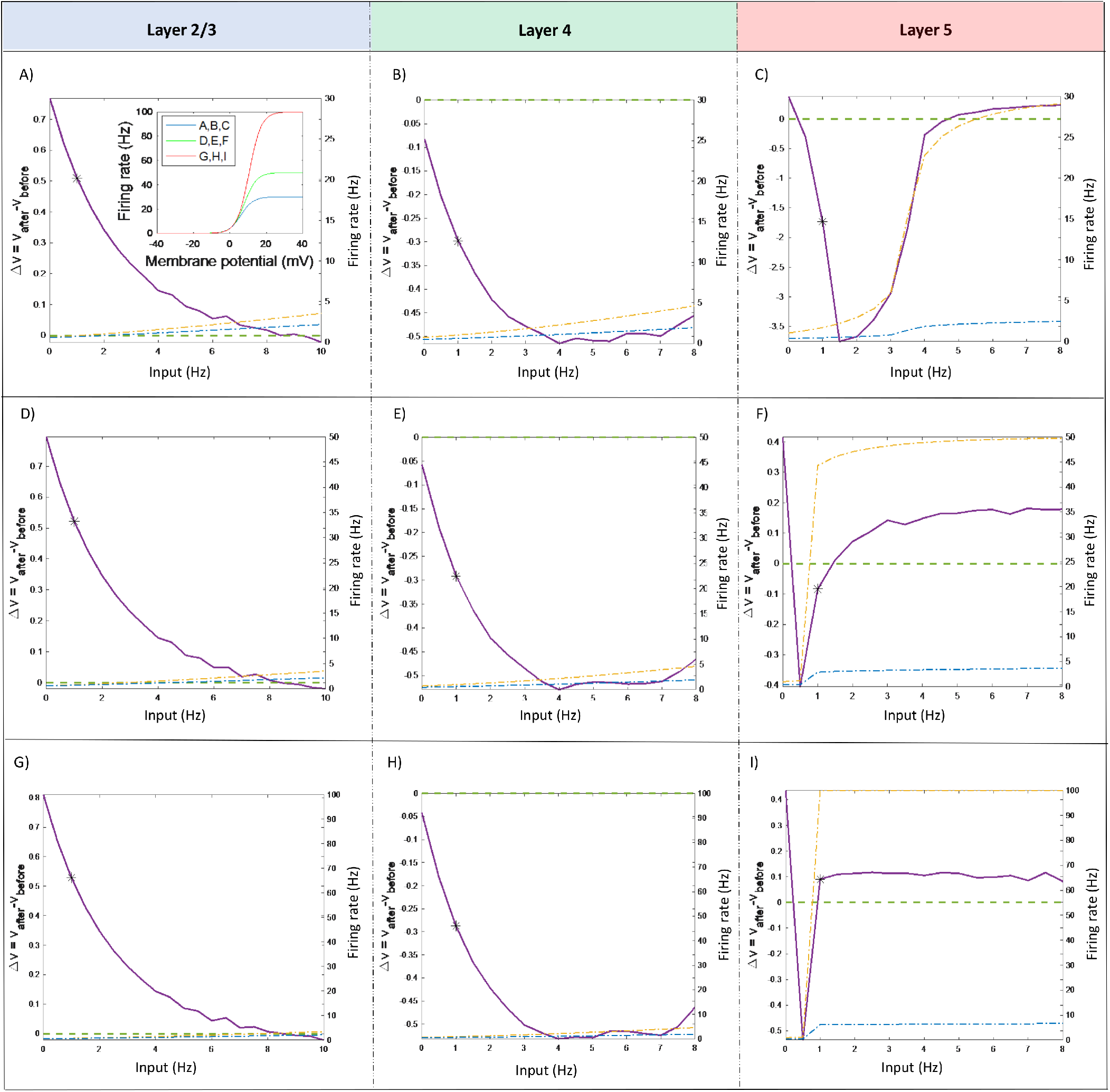
The effect of varying external input on the paradoxical effect in single motifs. A, B, C) The model of a single motif with the default sigmoid function (blue solid curve in the inset in panel A showing the relationship between membrane potential and firing rate). The purple curve indicates changes in the membrane potential (Δ*v*) of inhibitory neurons after perturbation with different input levels. The green dashed line indicates where Δ*v* is zero. Negative values of Δ*v* indicate paradoxical behaviour. The black star indicates the default value of the external input in the model (1 Hz). The orange dashed line shows the firing rate (right axis) of the excitatory population and the blue dashed line the firing rate of the inhibitory population. D, E, F) Effects of changing the external input on the paradoxical behaviour in the case where the sigmoid function has a maximum firing rate of 50 Hz (green solid curve in the inset in panel A). G, H, I) Effects of changing the external input on the paradoxical behaviour in the case where the sigmoid function has a maximum firing rate of 100 Hz (red solid curve in the inset in panel A).

To evaluate the effects of the parameters of the sigmoid function, we conducted the same experiment using higher maximum firing rates for the neurons. The different sigmoid function are shown in the panel inside Fig. 12A. The maximum firing rates were increased to 50 Hz and 100 Hz; respective thresholds were also adjusted to 8 mV and 10.5 mV in order to maintain the same input-output relationship below 10 Hz firing rate. The results in Fig. 12 show that the paradoxical behaviours of layer 2/3 (D and G) and layer 4 (E and H) were largely unaffected by the changes to the sigmoid function as the curves are similar to those in Fig. 12A and B, respectively. Layer 5 shows markedly different behaviour with higher maximum firing rate, with the excitatory neurons in layer 5 saturating at lower levels of external input, including at or close to the default value of 1 Hz.

## Discussion

### Inhibitory stabilization and paradoxical response

Inhibitory stabilization plays a key role in maintaining the excitatory-inhibitory balance, particularly in ISNs. The key question is whether ISNs and, consequently, inhibitory stabilization are present in the cortex. Neurophysiological recordings show a mixture of results displaying the presence of ISNs in the cortex [1, 3, 16, 17, 33]. In this study, we developed a neural mass model to investigate these differences across neurophysiological studies. The results indicate that there is a gradient of inhibitory stabilization across different layers of the cortex that depends on the level of excitation-inhibition balance in the network. The model shows that layer 5 of the cortex is more inhibitory stabilized and shows a strong paradoxical response to perturbation compared to layer 2/3 of the cortex, which does not show a paradoxical response (Fig. 5 and Fig. 10). These results accord with the studies that demonstrate the paradoxical effect in deeper layers of the cortex, such as layer 5, and studies that do not conclude that there is a paradoxical effect in the cortex based upon recordings from layer 2/3 of the cortex [3, 21].

This paradoxical response is the result of inhibitory-to-inhibitory connections that come into effect after a delay following a stimulus onset. Consequently, as the result of excitatory input to inhibitory neurons in ISNs, there is an initial transient increase in the membrane potential of inhibitory neurons that is followed by a sustained decrease in their membrane potential. To explore whether this decrease in the membrane potential of the neurons is the result of the recurrent inhibitory connections or inter-layer inhibitory connections, we investigated the paradoxical effect both in a single motif (Fig. 5) and the whole cortical column (Fig. 10). Both results show a paradoxical response in layer 4 and layer 5 and no paradoxical response in layer 2/3 of the cortical column. This finding indicates that the networks in different cortical layers are operating in the ISN regime regardless of inhibitory inter-layer connections. We have evaluated the robustness of the results by changing the level of the external input, connectivity constant and the nonlinear transfer function.

We have also investigated the ranges for different connectivity weights in which the network operates in the ISN regime and show the paradoxical response to perturbation (Fig. 6, Fig. 8, Fig. 7 and Fig. 9). The results indicate that the strength of the paradoxical response increases as the network is getting close to the bifurcation and reaches a maximum value just before the bifurcation. As a result of the saddle node bifurcation in the network, the firing rate of the excitatory neurons jump from low to high firing rate and saturate. This resembles a pathological condition such as an epileptic seizure. The key question arising from from these results is whether the strength of the paradoxical response can be used to predict the appearance of bifurcation in the network. As this bifurcation resembles a mathematical model of seizures in an epileptic brain, the question is whether measuring the strength of the paradoxical response over time can be used as an active biomarker to predict the occurrence of a seizure in the cortex, which remains to be explored in future studies.

### Excitation-inhibition balance

The default values for the connectivity weights across different populations in the model are based on the Synaptic Physiology database. This model shows that the variation in the connectivity strength provided by this database is not random and these variations in the connectivity strengths determine the gradual changes in the excitation-inhibition balance across different layers in the cortex. The model shows that the superficial layer 2/3 is more inhibitory dominant compared to deeper layers, such as layer 4 and layer 5. This also accords to neurophysiological and anatomical studies [22].

### The fixed-point analysis

The results of the fixed-point analyses show that increasing the excitatory-to-excitatory connectivity weight expands the regions where the fixed-point is bistable (Fig. 3A). However, an increase in the value of the inhibitory-to-inhibitory connectivity weight expands the region of having a stable fixed point in the network (Fig. 3A). On the other hand, the data in Table 1 shows that the lowest values for the connectivity weights are the recurrent excitatory-to-excitatory connections in each motif of a cortical column and the highest value for connectivity weights are recurrent inhibitory to inhibitory connections. This data also shows that in most cases the excitatory-to-inhibitory connections have higher connectivity weights than inhibitory-to-excitatory connections, which accords with the results obtained by the model that demonstrate a similar trend in the connectivity weights in order to be in a stable regime.

Increasing the excitatory-to-excitatory connectivity weight expands the regions where the neurons are either saturated or where the fixed-points are bistable. This is also the case for decreasing the value of the excitatory-to-inhibitory connectivity weight. Lower values of inhibitory-to-excitatory connectivity weight expand the regions where the fixed-point is stable but increases the likelihood of the saturation of the neurons. Therefore, the balance between excitation and inhibition in the network is the main factor that determines the region where the connectivity weights result in the operation of the network in a biologically plausible (non-pathological) regime.

### Assumption and approximation

Developing this model requires making a series of assumptions and approximations. Each unit in the model represent a population of neurons and the output of the model represents the average of the membrane potentials of the neurons in the population. Developing a detailed model of different types of neurons will shed light on several aspect of this study, which remain to be explored in future investigations. In this study, we have assumed the uniformity of the spike rates across each type of population of the neurons. The spike rate function is instantaneous and as it is only spiking rates that matter, which results in ignoring the variance in the spike rates of the neurons. The model of the synapses is current-based rather than conductance-based. A more realistic model would incorporate several biophysical features, such as dendritic filtering of synaptic input and drop-out of synapses at synaptic transmission. We also ignore activity-based effects upon the membrane time constants; the membrane time constants are always equal to the passive time constants. These issues will be explored in biopohysical models of the cortical neurons in future studies.

## Supporting information

**S1 Table.**
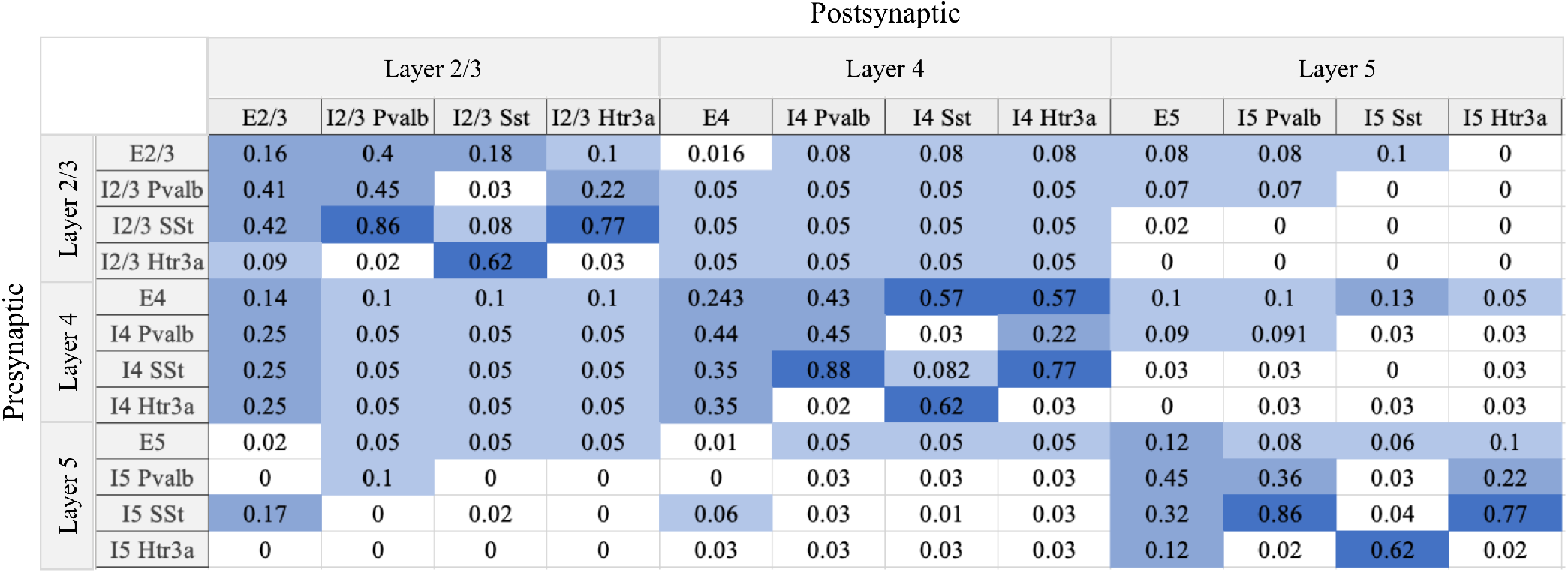
Connection probabilities. The connection probabilities across different types of neurons derived from the Figure 4A in [5]. E represents excitatory neurons and I represents inhibitory neurons. Inhibitory neuron classes are Htr3a, Sst and Pvalb. The highlighting indicates the level of connection probability, darker for higher chance of connection.

**S2 Table.**
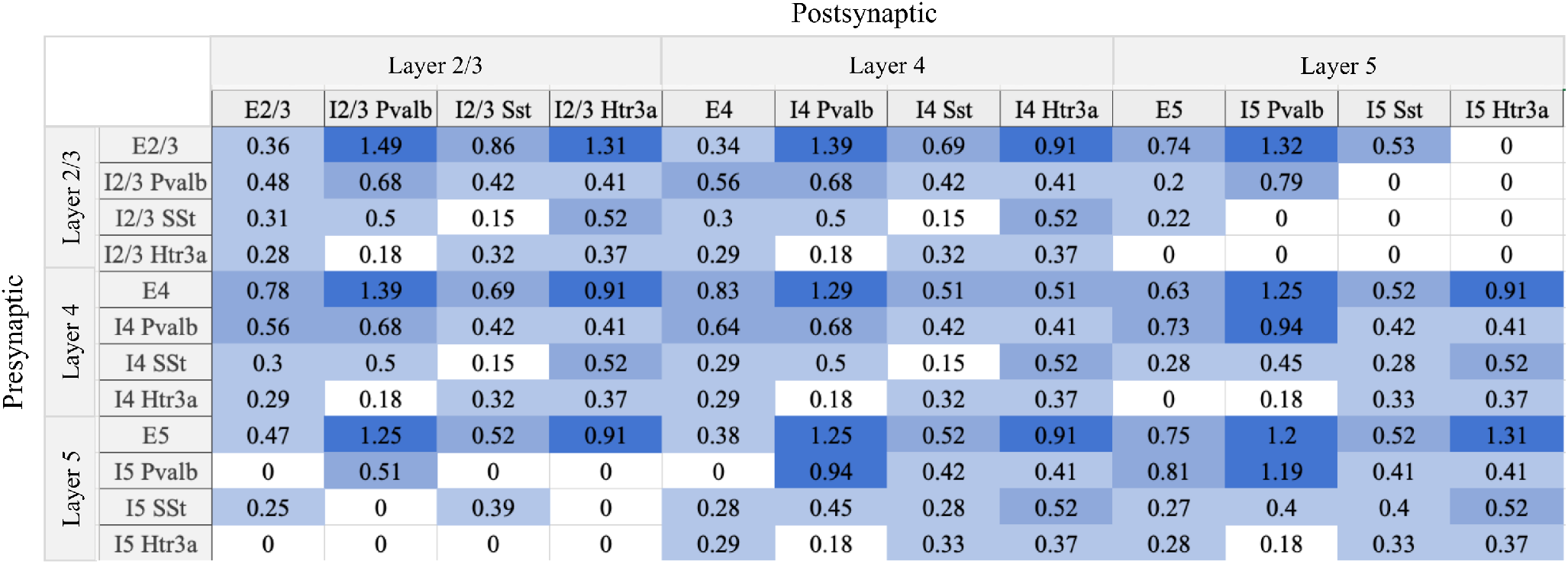
Synaptic strengths. The synaptic strengths between different types of neurons derived from the Figure 4B in [5]. E represents excitatory neurons and I represents inhibitory neurons. Inhibitory neuron classes are Htr3a, Sst and Pvalb. The highlighting indicates the strength of the synaptic connection, darker for stronger connections.

### SI 1. The equations describing the dynamics of membrane potentials and synaptic currents

The membrane potentials of the excitatory and inhibitory neural population in layer 2/3:

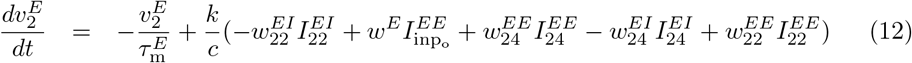

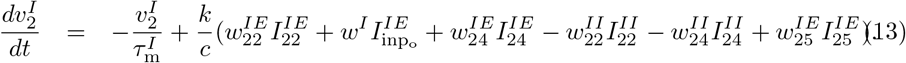

Synaptic currents from the excitatory population in layer 2/3 to other populations:

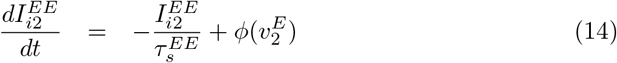

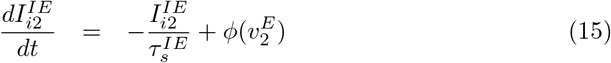

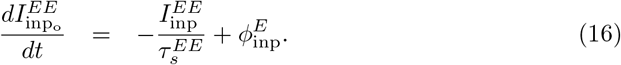

Synaptic currents from the inhibitory population in layer 2/3 to other populations:

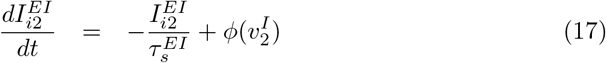

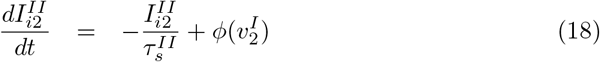

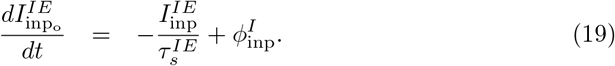

The membrane potentials of excitatory and inhibitory populations in layer 4:

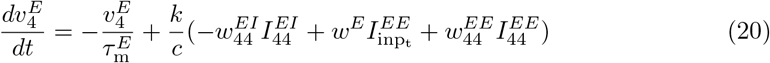

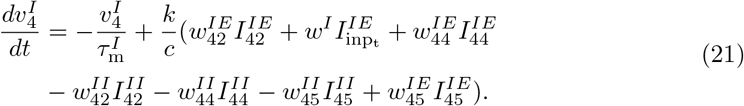

Synaptic currents from the excitatory population in layer 4 to other populations:

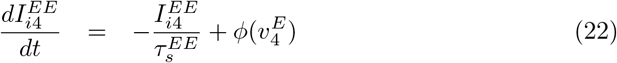

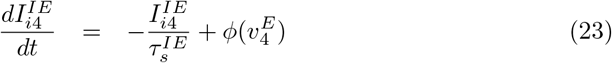

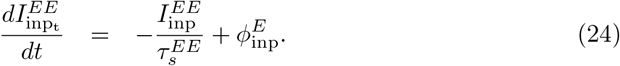

Synaptic currents from the inhibitory population in layer 4 to other populations:

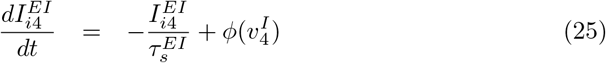

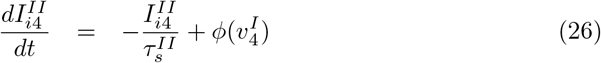

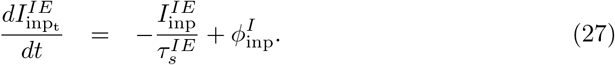

Membrane potentials of excitatory and inhibitory populations in layer 5:

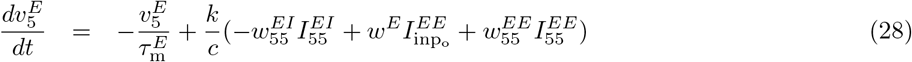

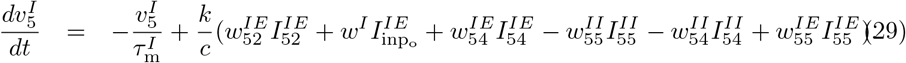

Synaptic currents from the excitatory population in layer 5 to other populations:

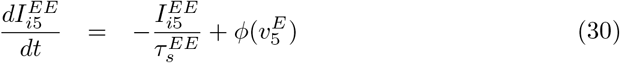

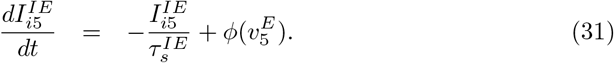

Synaptic currents from the inhibitory population in layer 5 to other populations:

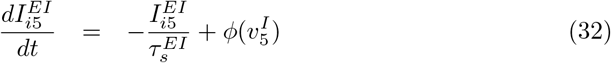

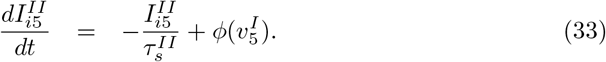

### SI 2. Calculation of fixed points

Setting the left-hand sides of equations Eq. (12)-Eq. (33) to zero results in the following equations for calculating the fixed points of the network:

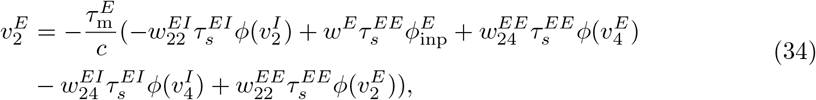

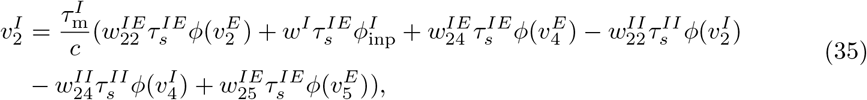

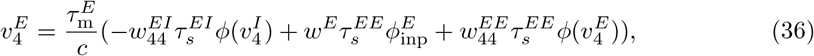

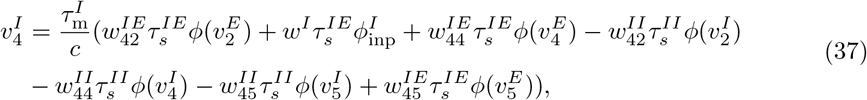

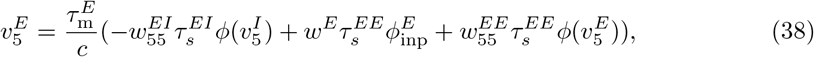

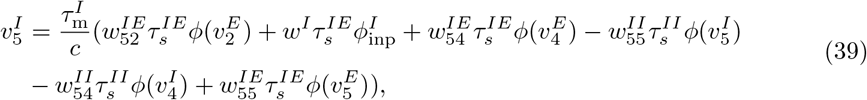

### SI 3. Variations in the excitatory and inhibitory nullclines by changing the connectivity weight

S1 Figure shows the variations in the location of the fixed point in the network (derived from intersection of inhibitory and excitatory nullclines) by changing the excitatory-to-excitatory connectivity weight in layer 2/3. The middle panel shows the variation in Δ*v* by changing the excitatory-to-excitatory connectivity weight. The surrounding panels shows the changes in the excitatory and inhibitory nullclines at different values of excitatory-to-excitatory connectivity weight indicated on the top of each panel. The black arrows indicate the associated value of Δ*v* for each panel. Top-left panel shows the excitatory and inhibitory nullcline when the excitatory-to-excitatory connectivity weight equals to the default value of 0.06. The intersection of excitatory and inhibitory nullcline is on the negative slope of the excitatory nullcline. The intersection of the inhibitory nullcline after perturbation and excitatory nullcline moves to a higher value of membrane potential of inhibitory neurons indicating a non-paradoxical effect. Increasing the strength of excitatory-to-excitatory connectivity weight results in gradual shift of the intersection point to positive slope where the network shows a paradoxical effect until the connectivity weight equals to 0.249. Further increase in the value of the connectivity weight shifts the intersection to a negative slope of the excitatory nullcline where there is no paradoxical effect.

**S1 Figure.**
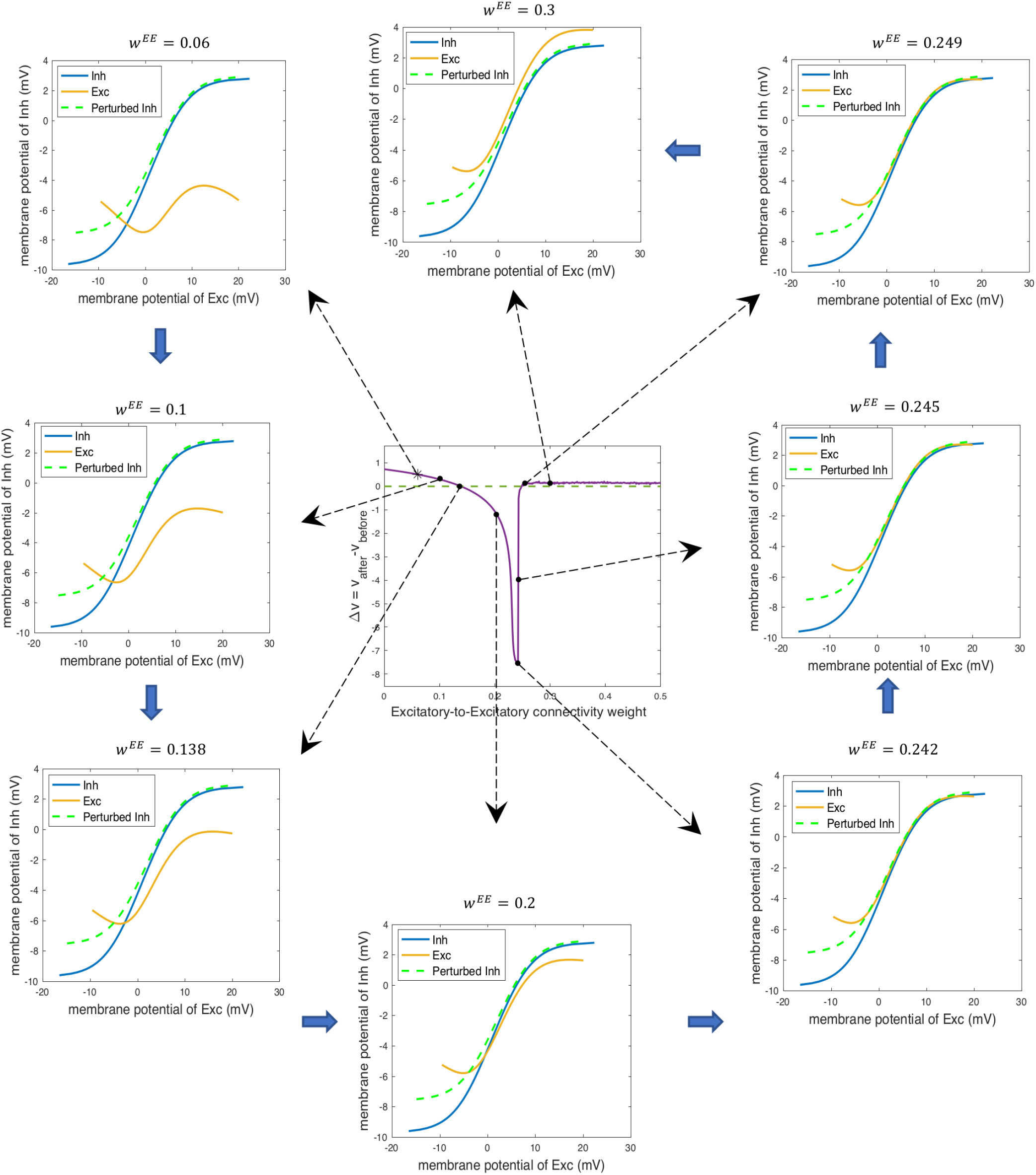
The variations in the location of the fixed point by changing the excitatory-to-excitatory connectivity weight. The purple curve in the middle panel shows the variation in Δ*v* by changing excitatory-to-excitatory connectivity weight. The surrounding panels show the variations in the excitatory and inhibitory nullclines of a single motif by changing the excitatory-to-excitatory connectivity weight. The value of the excitatory-to-excitatory connectivity weight is given at the top of the panel. The black arrows indicate the panel related to the specific values of Δ*v* at that connectivity weight. The orange curve in the surrounding panel shows the excitatory nullcline, the blue curve is the inhibitory nullcline, and the green dashed curve shows the inhibitory nullcline after perturbation.

## Acknowledgments

This work was funded by the Australian Government, under the Australian Research Council’s Training Centre in Cognitive Computing for Medical Technologies (project number ICI70200030).

## Footnotes

1 The Synaptic Physiology database is publicly available at https://portal.brain-map.org/explore/models/mv1-all-layers

2 https://sourceforge.net/projects/matcont/

